# An integrated multi-tissue atlas of epigenomic landscapes and regulatory elements in the bovine genome

**DOI:** 10.1101/2025.08.21.671512

**Authors:** Dailu Guan, Jennifer Jessica Bruscadin, Wenjing Yang, Claire Prowse-Wilkins, Junjian Wang, Bruna Petry, Houcheng Li, Huicong Zhang, Xiaoqin (Shally) Xu, Ying Wang, Zhangyuan Pan, Yuri Utsunomiya, Gordon K. Murdoch, Kimberly M. Davenport, Yue Xing, Guosong Wang, Christian Maltecca, Li Ma, Gonzalo Rincon, Sarah Corum, Pengcheng Lyu, Nader Deeb, Virgínia Mara Pereira Ribeiro, Jennifer J. Michal, Zhiping Weng, Jicai Jiang, Lingzhao Fang, Honglin Jiang, Brenda M. Murdoch, Monique Rijnkels, Clare A. Gill, Timothy P. L. Smith, Zhihua Jiang, Wansheng Liu, James Reecy, Juan F. Mendrano, James E. Koltes, Pablo J. Ross, Huaijun Zhou

**Author notes:** **Corresponding authors:** Prof. Huaijun Zhou: Department of Animal Science, University of California-Davis, Davis, CA, 95616, USA. These authors contributed equally (Dailu Guan, Jennifer Jessica Bruscadin, Wenjing Yang, Claire Prowse-Wilkins). **Emails for all authors:**;,;,;,;,;.

## Abstract

Deciphering the regulatory syntax of the genome is essential to understand the genetic and molecular architecture of complex traits, as most trait-associated variants lie in non-coding regions. Yet, functional annotation of the bovine genome remains limited, hindering our ability to unravel the mechanisms underpinning complex traits of economic and ecological importance in cattle. Here, we present a comprehensive epigenetic atlas comprising 1,138 genome-wide epigenetic profiles, including chromatin accessibility, six histone modifications, CCCTC-binding factor (CTCF) transcription factor binding, DNA methylation, chromatin conformation, and transcriptomes across 53 adult tissues, five fetal tissues, and seven primary cell types. This atlas-level data enables us to annotate around 45% of the genome as putative regulatory elements exhibiting tissue- or cell-specific regulatory activity. Leveraging sequence-to-function deep learning models, we discovered 301 sequence motifs and predicted the functional impact of genetic variants through *in silico* mutagenesis, thereby facilitating the decoding of the regulatory syntax of the cattle genome and fine-mapping of GWAS loci for 22 complex traits. Cross-species analysis further revealed evolutionarily conserved features of regulatory architecture and provided evolutionary insights into complex traits and diseases in humans. Together, this atlas offers a foundational resource for advancing cattle functional genomics, sustainable breeding, and studies of regulatory evolution.

## Introduction

Cattle are a key agricultural species, providing meat, milk, leather, and labor, while also serving as a valuable model to study embryo development, epigenetics and imprinting, immunology, and the genetics of complex traits^1,2^. Advances in high-throughput sequencing and genotyping technologies have substantially enhanced our understanding of the genetic architecture of economically important traits in cattle and accelerated the pace of genetic improvement^3^. For instance, incorporation of single nucleotide polymorphisms (SNPs) into dairy breeding programs has doubled the rate of genetic gain for key production traits^4^. As of the latest update, the CattleQTLdb^5^ catalogs 193,453 quantitative trait loci (QTL), the majority of which reside in non-coding genomic regions^6^. This strongly suggests that many QTL act through regulatory mechanisms, modulating transcriptional activity of nearby or distal gene regulatory elements^7^. Comprehensive annotation of the non-coding bovine genome is therefore essential to uncover mechanistic insights into genome-wide association study (GWAS) signals and pinpoint causal variants. Such insights may ultimately enable precision breeding strategies and the development of “ultimate genotypes”^8^ through genome editing^9^ and synthetic biology^10^, while further enhancing the utility of the cattle as a biomedical model.

Initial efforts to characterize bovine regulatory landscape have provided important foundations. For example, Kern et al.^11^ performed genome-wide annotation of regulatory elements by analyzing chromatin immunoprecipitation followed by sequencing (ChIP-seq) for four core histone modifications (H3K4me3, H3K27ac, H3K4me1, H3K27me3) and the DNA-binding protein CTCF, as well as Assay for Transposase-Accessible Chromatin using sequencing (ATAC-seq), across eight core tissues (cortex, cerebellum, hypothalamus, liver, lung, spleen, muscle, adipose). Subsequent studies extended these annotations to rumen tissues^12,13^ and rumen epithelial primary cells^12^, and incorporated additional assays such as Cap Analysis Gene Expression (CAGE) sequencing^14^, as well as broader genomic feature mapping^15^. Although these studies parallel large-scale efforts like ENCODE^16^ and Roadmap Epigenomics^17^ in humans towards annotating the non-coding genome, the bovine non-coding genome remains incompletely annotated. A systematic, scalable effort spanning diverse tissues, cell types, and developmental stages (hereafter referred to as “tissue contexts”^18^) is essential to fully characterize context-specific regulatory mechanisms.

Complementary transcriptomic resources, including the Cattle Genotype-Tissue Expression (CattleGTEx)^19–21^ and large-scale allele-specific (AS) analysis^22^, have identified millions of regulatory variants influencing gene expression and splicing. Yet, these resources have not yet advanced to systematic elucidation of causal regulatory mechanisms underlying GWAS associations of complex traits in cattle. Meanwhile, the emergence of sequence-to-function (S2F) deep learning models offers powerful approaches to decode the regulatory syntax of the genome and to prioritize functional variants^23^. Tools like SpliceAI^24^ and Pangolin^25^ have demonstrated utility in predicting splicing effects in cattle^26^, but broader application of S2F models to interpret regulatory elements and context-specific variants remains largely unexplored.

In this study, we constructed a comprehensive bovine epigenomic atlas by integrating 540 newly generated datasets from 25 previously unprofiled tissues and assays with public available data^12,13,15,27,28^. In total, our resource comprises 1,138 genome-wide epigenomic profiles including 158 RNA-seq, 202 ATAC-seq, 91 whole genome bisulfite sequencing (WGBS), 682 ChIP-seq (six histone marks and CTCF), and 5 chromatin conformation (Hi-C) datasets. We used this data atlas to annotate chromatin states, link enhancers to their target genes, decode regulatory syntax via deep learning approaches, fine-map GWAS loci, and perform multi-tissue cross-species comparisons (**Fig. 1a**). The resulting resource provides a powerful foundation for mechanistic dissection of complex traits, prioritization of functional variants, and comparative regulatory biology.

**Figure 1.**
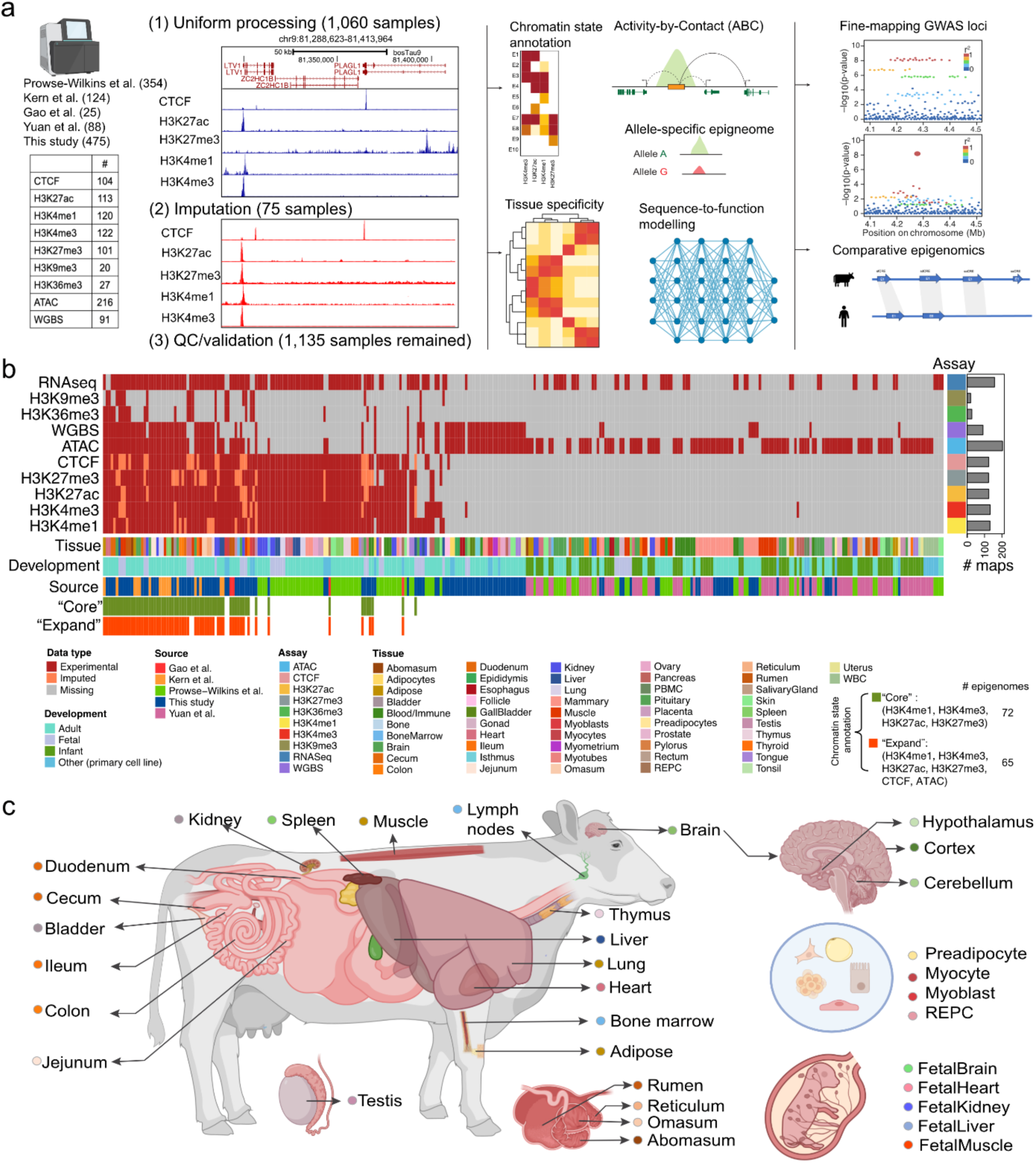
Overview of the study design and datasets. (a) Schematic overview of the analytical framework used in this study, including data processing, imputation of missing or low-quality histone marks, chromatin state annotation, enhancer-gene linking, tissue-specificity analysis, sequence syntax modelling using deep learning, fine-mapping GWAS loci of complex traits, and cross-species epigenomic comparison. (b) Summary of multi-omic datasets included in this study. (c) Illustration of tissues used for chromatin state annotation in this study. All samples for these tissues have either observed or imputed all four core histones (i.e. H3K4me1, H3K4me3, H3K27ac, and H3K27me3). Icons and the cattle image were adapted from BioRender.com.

## Results

### A comprehensive atlas of the bovine epigenomic landscape

We have compiled a comprehensive atlas of 1,138 genome-wide epigenomic maps through uniform processing and quality control (**Fig. 1a-b**, **Supplementary Figures S1-S3**, **Table S1-S4**, see **Methods**). These include 158 RNA-seq maps to quantify gene expression (**Supplementary Figure S4, Table S1**), 202 ATAC-seq datasets to profile open chromatin regions (OCRs) (**Supplementary Figure S5-S6**, **Table S2**), 91 WGBS datasets to measure DNA methylation (**Supplementary Figure S7-S8**, **Table S3**), 682 ChIP-seq datasets capturing histone modifications and transcription factor (TF) binding (**Table S4**), and 5 public Hi-C datasets (**Table S5**). The ChIP-seq datasets encompass H3K4me1 (n = 131), marking enhancer regions; H3K4me3 (n = 132), associated with promoter activity; H3K27ac (n = 125), indicative of active enhancer and promoter regions; H3K27me3 (n = 123), marking Polycomb-mediated repression; H3K9me3 (n = 20), linked to heterochromatin; H3K36me3 (n = 27), associated with actively transcribed regions; and CTCF (n = 124), marking chromatin organizing elements^29^ (**Supplementary Figure S9**). Among these, 218 ChIP-seq datasets (32%) from 25 tissues were newly generated in this study, 389 were retrieved from public data^12,13,15,28^, and 75 datasets were computationally imputed using the ChromImpute pipeline^30^. They include H3K4me1 (n = 11), H3K4me3 (n = 10), H3K27ac (n = 12), H3K27me3 (n = 22), and CTCF (n = 20), with genome-wide Pearson correlation ranging from 0.56 (H3K27me3) to 0.91 (H3K4me3), comparable to those observed in human datasets^30^ (**Extended data Fig. 1, Supplementary Figure S10-S11**). Peak calling from experimental ChIP-seq datasets yielded 895,108 H3K4me1 (52.7% of the autosomal genome), 737,720 H3K4me3 (18.2%), 802,966 H3K27ac (26.9%), 1,038,083 H3K27me3 (47.8%), 442,773 H3K9me3 (7.2%), 567,480 H3K36me3 (18.4%), 831,902 CTCF (19.8%) non-overlapping peaks. ATAC-seq analysis identified 953,179 non-overlapping open chromatin regions (OCRs) covering 21% of the genome, while WGBS profiling identified approximately 1.8 million methylation blocks (i.e. regions with ≥4 continuous CpGs), covering 73.4% of the genome. Additionally, uniform processing Hi-C datasets yielded on average 4,979 topologically associating domains (TADs) per sample at 10 kb resolution (**Table S5**). Altogether, we provide a comprehensive resource to explore the regulatory landscape of the cattle genome.

### Comprehensive regulatory element annotation of the bovine genome

Leveraging the atlas-level maps above, we defined a 10-state “core” chromatin model across 72 bovine epigenomes representing 33 tissues (**Fig. 1c**) (**Fig. 2a**, **Supplementary Figure S12**). Here, an epigenome was defined as a biosample with all four core histone marks (i.e., H3K4me1, H3K4me3, H3K27ac, and H3K27me3). These states^31–33^ included three promoter-like states (PLS): TssA (active transcription start site [TSS], covering 2.2% of the genome across tissue contexts), TssFlnk (flanking TSS, 3%), and TssWk (weak flanking TSS, 3.5%); three enhancer-like states (ELS): EnhA (active enhancer, 7.6%), EnhWk (weak enhancer, 11.7%), and EnhPos (poised enhancer, 12.5%); two bivalent states: BivTss (bivalent TSS, 0.98%) and BivEnh (bivalent enhancer, 0.68%); as well as ReprPC (repressed Polycomb, 4.6%) and Quies (quiescent/low signal) **(Fig. 2a)**. These chromatin states exhibited distinct patterns of gene expression (**Fig. 2a**), chromatin accessibility (**Extended data Fig. 2a**), DNA methylation (**Extended data Fig. 2b**), evolutionary conservation (**Fig. 2a, Extended data Fig. 2c, Supplementary Figure S13**), and transposable elements enrichment (**Supplementary Figure S14-15**). Of note, we found ELS from fetal epigenomes were significantly enriched for CR1 and L2 subfamilies (**Supplementary Figure S15**), and a high number of ELS overlapped with LINE (e.g. L1 and L2) and SINE subfamilies (**Supplementary Figure S16**).

**Figure 2.**
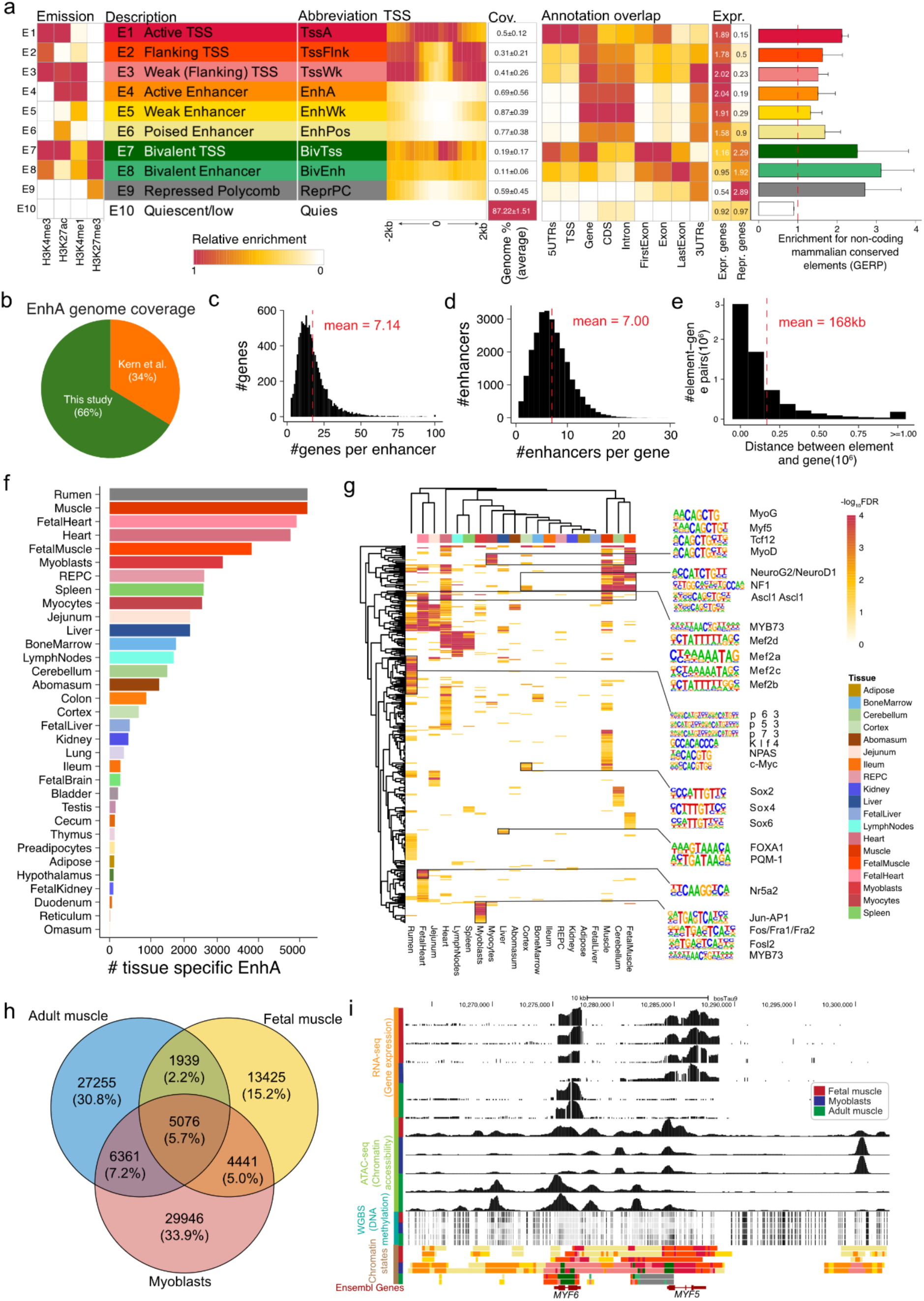
Chromatin state annotation and context-specific regulatory dynamics. (a) Definition and characterization of chromatin states, including emission probabilities, functional descriptions (annotations), abbreviations, enrichment near transcription start sites (TSS), overlap with genomic features, associated gene expression levels, and evolutionary constraint based on GERP scores^101^ (Ensembl release v105). CDS: coding sequence. UTR: untranslated regions. (b) Genome coverage of strong enhancers (EnhA) compared to annotations from that of Kern et al.^11^ (c)-(e) Enhancer-gene linking based on the Activity-by-Contact (ABC) model^34^: number of genes linked per enhancer (c), number of enhancers linked per gene (d), and genomic distance between enhancers and their predicted target genes (e). (f) Number of tissue-specific strong enhancers (EnhA) identified across tissues. (g) Motif enrichment analysis of tissue-specific EnhA elements. Representative enriched motifs and associated transcription factors (TFs) with known tissue relevance are shown at right. (h) Venn diagram depicting the overlap of strong enhancers (EnhA) identified in myoblasts (cell line), adult muscle and fetal muscle (bulk tissue). (i) Genome browser tracks displaying gene expression and epigenomic signals around the *MYH5* and *MYH6* loci (chr5:10262831-10302829 bp).

Compared to the previous annotation of eight core tissues by Kern et al.^11^, our atlas incorporates 25 additional tissues (16 adult, 5 fetal and 4 primary cell types), resulting in an additional 106 Mb of annotated sequences (19% increase). For example, we annotated 146,445 non-overlapping EnhA, of which 66% were novel^11^ (**Fig. 2b**). To expand annotation, we further constructed a 15-state “expanded” model across 65 epigenomes that included chromatin accessibility and CTCF data. This model introduced additional states such as TssFlnD (downstream flanking TSS), EnhAHet (active enhancer lacking ATAC signal), EnhCtcf (enhancer with CTCF), CtcfIsl (CTCF island), AtacIsl (ATAC island), and BivTssCtcf (bivalent TSS with CTCF) (**Extended data Fig. 3**).

To link enhancers to their target genes, we employed the Activity-by-Contact (ABC) approach^34^, and successfully linked 25,567 unique enhancers to 26,100 genes, resulting in a total of 182,596 enhancer-gene pairs (**Fig. 2c-e, Supplementary Figure S17**). On average, each enhancer was associated with 7.1 genes (**Fig. 2c**), and each gene with 7 enhancers (**Fig. 2d**), the mean enhancer– gene distance was 168 kb (**Fig. 2e**). Furthermore, using ROSS software^35^, we predicted an average of 1,242 super-enhancers across 32 tissue contexts (covering 18.6% of the genome), ranging from 6 in liver to 3,128 in cerebral cortex (**Supplementary Figure S18-19**). Altogether, approximately 45% of the bovine autosomal genome (excluding Quies) was annotated as regulatory elements, representing the most comprehensive regulatory annotation to date in cattle.

### Dynamic epigenomic signatures across tissues and cell types

To investigate context-specific regulation of gene expression, we systematically analyzed chromatin state dynamics across tissues, developmental stages, and primary cell types. Tissue-specific states were first identified (**Supplementary Figure S20** and **Table S6)**. Active enhancer (EnhA) showed the strongest tissue specificity, with muscle tissues, including muscle, heart, fetal heart, fetal muscle, and myoblasts, exhibiting the highest number of tissue-specific EnhA elements (**Table S6**). In contrast, digestive tissues such as omasum, reticulum, and duodenum (excluding rumen) showed relatively few unique EnhA elements (**Fig. 2f**). These tissue-specific EnhA elements were strongly enriched for biologically relevant functional terms (**Supplementary Figure S21, Table S7**) and TF motifs^36^ (**Fig. 2g, Table S8**). For instance, motifs for MYOG, MYF5, and MYOD were enriched in muscle tissues, NEUROG2, NEUROD1, and NF1 in central nervous system (CNS) tissues, JUN, FOS and FRA1 in myoblasts (**Fig. 2g**), consistent with their established roles in cell proliferation and differentiation^37,38^. To illustrate, we compared muscle-related tissue contexts, including adult and fetal bulk muscle, as well as myoblasts (**Supplementary Figure S22-24, Table S9-S10**). As shown in **Fig. 2h**, we identified 27,255 adult-specific (30%), 13,425 fetal-specific (15%), and 29,946 myoblast-specific (34%) EnhA elements.

A representative example involves the myogenic regulators *MYF5* and *MYF6*. While *MYF6* is highly expressed in adult and fetal muscle, but not in myoblasts, *MYF5* shows high expressions in fetal muscle and myoblasts, but not in adult muscle (**Fig. 2i, Supplementary Figure S24**). Chromatin accessibility data mirrored these expression patterns, with myoblasts- and fetal muscle-specific enhancers flanking the *MYF5* locus (**Fig. 2i**). These findings align with the established role of *MYF5* in early myogenic differentiation and *MYF6* in terminal myogenic differentiation^39^, mediated by a combinatorial network of enhancer elements during development^40^.

We next explored DNA methylation dynamics using 91 newly generated WGBS datasets (**Supplementary Figure S7-S8**), identifying 207,436 differentially methylated blocks (DMBs) (see **Methods**, **Table S11-S12**) across approximately 1.8 million tested regions (**Table S11**). To construct a bovine tissue-specific hypomethylated atlas, we selected the top 50 differentially unmethylated regions per tissue (for tissues with >50 DMBs), resulting in a total of 1,120 DMBs (**Extended Data Fig. 4**). These DMBs exhibited low methylation level (average 18%) and strong co-enrichment with chromatin accessibility and histone modifications, as exemplified in the cerebellum (**Extended Data Fig. 5a**). Motif enrichment analysis using HOMER^36^ further revealed tissue-relevant TFs (**Table S13**, **Extended Data Fig. 5b**), such as NEUROD1 in cerebellum, MYB81 in muscle, and MYF5 in myoblast (**Extended Data Fig. 5b**). Notably, *MYF5* encodes a critical regulator of skeletal myogenesis^41^. The tissue-specific patterns for chromatin accessibility are shown in **Supplementary Fig. 25**, and for gene expression in **Supplementary Fig. 26**. Finally, integration of tissue-specific OCRs with GWAS data revealed significant enrichment of complex trait loci in context-matched regulatory landscapes (**Extended Data Fig. 6**), underscoring the functional relevance of these dynamic epigenomic signatures.

### Deep learning reveals regulatory sequence syntax across tissue contexts

Gene expression is initiated and regulated through the binding of TF to specific DNA sequence, known as TF binding site (TFBS)^42^. Recent advances in S2F deep learning models have demonstrated strong performance in decoding regulatory syntax of DNA sequences and predicting TFBS. To explore regulatory syntax of the cattle genome, we trained ChromBPNet^43^, a convolutional neural network (CNN), to model the relationship between genomic sequence and chromatin accessibility across 55 tissue contexts. For each tissue, model performance, evaluated as the Pearson correlation between predicted and observed log-transformed ATAC-seq read counts on held-out peaks (**Methods**), achieved a median of 0.76 (**Extended Data Fig. 7a, Table S14**), comparable to reports in human datasets^43,44^. Importantly, Pearson correlations were stable across distances to the nearest TSS (**Extended Data Fig. 7b**). To assess robustness, we applied the models to independent ATAC-seq data from Yuan et al.^27^ across 19 overlapping tissues. For example, in abomasum, our model achieved a Pearson correlation of 0.84 on our datasets (**Extended Data Fig. 7c**) and 0.75 on Yuan et al.’s dataset^27^ (**Extended Data Fig. 7d**), with differences potentially reflecting breed or age effects^27^. A representative locus at *DGAT1* (chromosome 14: 603,067-604,765) across tissues in both datasets, achieved a Pearson correlation of 0.69, with bone marrow showing the highest chromatin accessibility (**Extended Data Fig. 7e**). Additionally, the model could also predict tissue-specific OCR peaks, as demonstrated in **Extended Data Fig. 7f**.

We next used high-performing models (Pearson’s *r* ≥ 0.72, 44 tissues retained) for *de novo* motif discovery (**Extended Data Fig. 7a**). Using DeepLIFT^45^ to score nucleotide contribution and TF-MoDISco^46^ to cluster predictive subsequences, we identified a non-redundant set of 301 *de novo* motifs across 44 selected tissues (average: 16 per tissue, range: 9–23; **Fig. 3a**). The canonical CTCF motif was consistently recovered in all tissue contexts, with contribution weight matrices (CWM) strongly resembling the JASPAR positional weight matrix (PWM) (**Fig. 3b**), and closely matching motifs identified in humans^44^. We also recovered some well-known TF motifs (e.g., NEUROD1 in cerebellum and MEF2B in heart) (**Fig. 3c**) as well as motifs with limited similarity to entries in the JASPAR database (*q*-value < 0.05), suggesting under-characterized or potentially novel TFBSs (**Fig. 3d**). Motif scanning across all OCRs revealed KLF15, a zinc-finger transcription factor with key roles in metabolic processes^47^, as the most frequently occurring motif across tissues (**Fig. 3e**). Collectively, our findings demonstrate that ChromBPNet generalizes well to cattle, enabling accurate prediction of chromatin accessibility and discovery of conserved as well as novel sequence motifs. This supports its utility in decoding *cis*-regulatory logic in non-human species.

**Figure 3.**
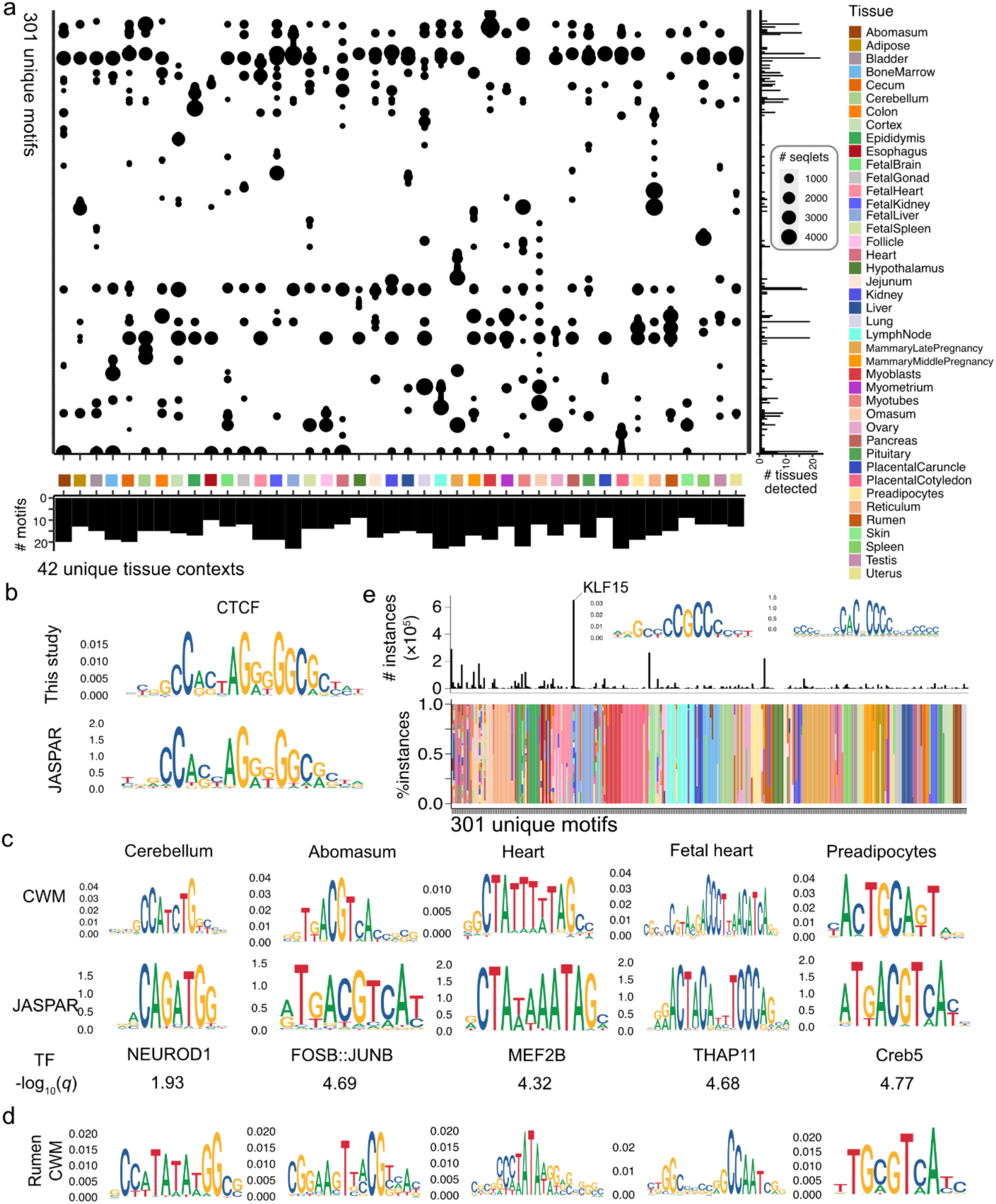
Sequence motif discovery across tissue contexts. (a) Number of seqlets (i.e., short informative subsequences) supporting each of the 301 unique motifs (y-axis) across 42 tissue contexts (x-axis). (b) Comparison between the CTCF motif identified by ChromBPNet^43^ in this study (top) and the canonical CTCF motif from the JASPAR database (bottom). (c) Representative motifs identified in this study showing high similarity to known TF motifs curated in the JASPAR database. (d) Five motifs discovered in the rumen, a ruminant-specific organ, which show low similarity to any known motifs in JASPAR motifs (*q*-value > 0.05). (e) Proportion of motif instances identified across tissue contexts. The KLF15 motif was the most frequently detected. The unit of y-axis for motifs patterns is “bits”, while the ChromBPNet derived motifs are represented using contribution weight matrix (CWM) score. The known non-redundant motifs from the JASPAR database were last accessed on February 11, 2022. TF: transcription factor.

### An atlas of allele-specific regulatory variations

To assess the impact of genetic variants on regulatory activity, we performed allele-specific (AS) analyses across about 10 million SNPs identified from two individuals (see **Methods**). Out of 3,568,140 heterozygous SNPs overlapping epigenomic peaks, 91,125 (2.55%) exhibited significant allelic imbalance (adjusted *P*^48^ < 0.05, **Fig. 4a, Table S15**). AS SNPs were enriched in gene-dense and regulatory regions, showing positive correlation with gene density (Pearson’s *r* = 0.23, *P* = 3.92×10^-32^), ChIP-seq peak density (Pearson’s *r* = 0.15, *P* = 8.7×10^-15^) and OCRs (Pearson’s *r* = 0.18, *P* = 3.04×10^-20^) (**Fig. 4a**). The majority (70%) of AS SNPs were located in intronic or intergenic regions (**Fig. 4b**), with significant enrichment within 5 kb upstream of TSS, particularly promoter marker H3K4me3 (**Fig. 4c**). AS effects on gene expression correlated strongly with those affecting active epigenomic marks (**Fig. 4d**), highlighting coordinated regulatory roles. Effect sizes of RNA-seq AS SNPs were highly and significantly correlated with those previously reported by Prowse-Wilkins et al.^49^(**Fig. 4e**), though they showed limited concordance with eQTLs from the multi-breed cattle GTEx dataset^19,50^ (**Supplementary Figure S27a**). Furthermore, effect sizes of epigenomic AS SNPs identified in our analysis showed high concordance with the Prowse-Wilkins dataset^49^ (**Fig. 4f**).

**Figure 4.**
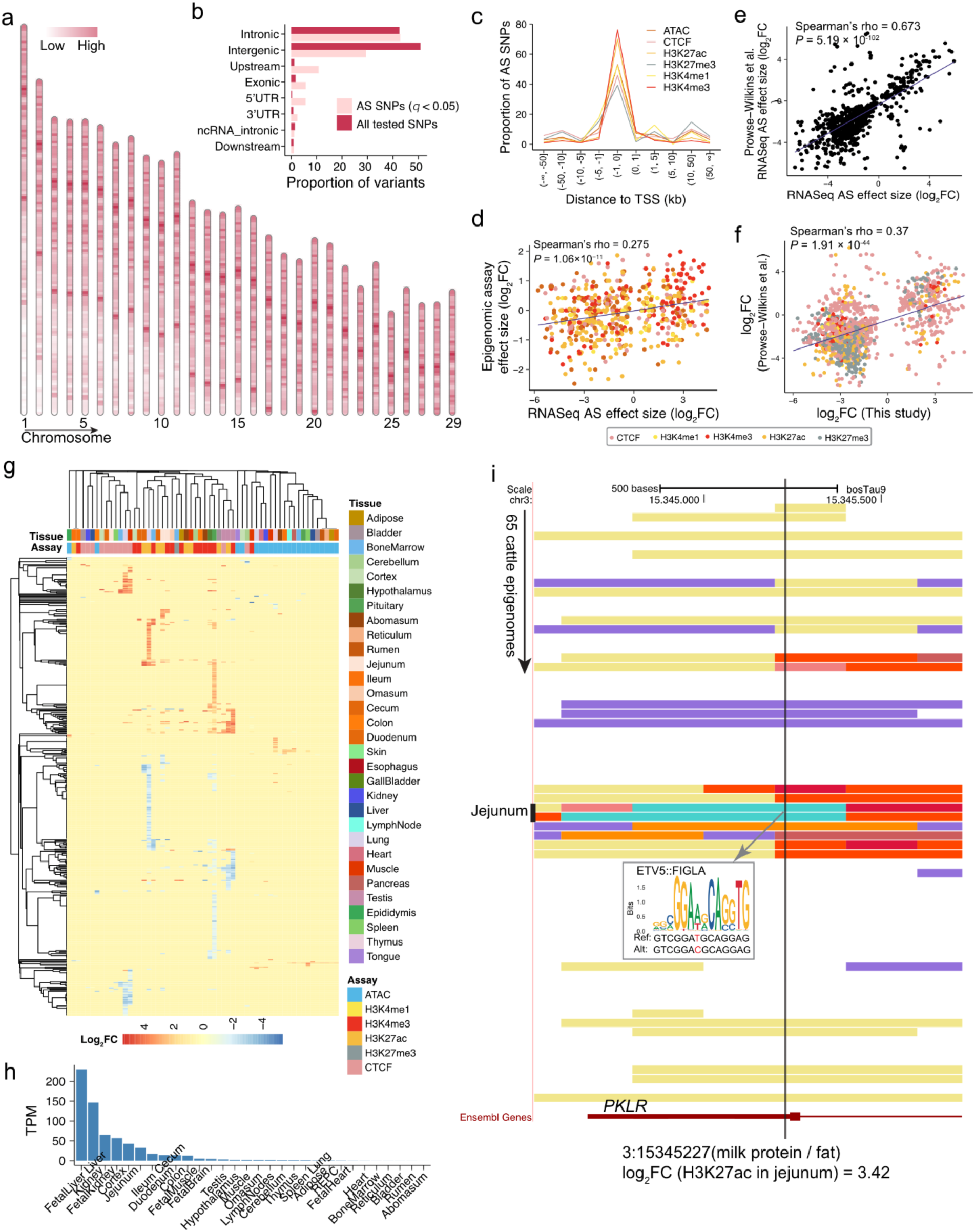
Identification and characterization of allele-specific (AS) epigenomic variation. (a) Genome-wide distribution of AS SNPs across bovine autosomes. (b) Proportion of SNPs (x-axis) by variant categories (y-axis). The variant category was obtained using SnpEff^102^. UTR: untranslated region. (c) Genomic distribution of significant AS SNPs relative to gene transcription start sites (TSS). (d) Correlation of AS effect sizes (log_2_ scaled) in RNA-seq (x-axis) and matched epigenomic assays (y-axis). (e) Comparison of AS effect sizes (log_2_ scale) in RNA-seq between this study (x-axis) and Prowse-Wilkins et al.^49^ (y-axis). (f) Comparison of AS effect sizes (log_2_ scale) in epigenomic assays between this study (x-axis) and Prowse-Wilkins et al.^49^ (y-axis). *P*-values in (d–f) were computed using the asymptotic *t-*distribution approximation. FC: fold change. (g) Heatmap showing fold changes (log_2_ scale) for 439 AS SNPs overlapping known trait-associated QTLs curated in the CattleQTLdb^5^. (h) Gene expression profile of *PKLR* across tissues measured in transcripts per million (TPM). (i) Genomic tracks showing an AS SNP in the promoter region of *PKLR*, detected in jejunum H3K27ac data. This SNP overlaps a predicted ETV5:FIGLA binding motif (identified using FIMO^103^).

To evaluate their functional relevance to complex traits, we intersected AS SNPs with trait-associated SNPs curated in the CattleQTLdb^5^, identifying 439 putative variants (**Supplementary Figure S27b, Table S16**). These AS SNPs exhibited strong assay- and tissue-specific epigenomic effects (**Fig. 4g**). For example, SNP 3:15345227 in the *PKLR* promoter, a gene associated with milk traits^51^ and highly expressed in brain-related tissues, exhibits a strong H3K27ac AS signal in jejunum (log_2_FC = 3.42), coinciding with a tissue-specific BivTssCtcf state (**Fig. 4h-i**). Another variant, SNP 29:42051788 in the *RTN3* promoter, showed the largest AS fold change in muscle, affecting H3K4me3 signal (**Supplementary Figure S27c-d**). *RTN3* has been associated with tenderness score^52^ and lipid metabolism^53^, suggesting potential causal functionality.

Together, these findings establish an atlas of allele-specific regulatory variants in cattle, linking genetic variation to tissue-specific regulatory effects and providing candidate functional variants underlying complex traits.

### *In silico* prioritization of regulatory variants

S2F modeling enables quantitative prediction of the regulatory impact of genetic variants through *in silico* mutagenesis. To systematically prioritize functional variants across tissue contexts, we employed deltaSVM^54^, which balances context coverage (e.g. tissue and assay types) with computational efficiency (**Extended Data Fig. 8**). Models were trained using epigenomic peaks as positives and GC-matched controls as negatives across 206 assay–tissue combinations (55 OCR, 29 CTCF, 32 H3K27ac, 27 H3K27me3, 30 H3K4me1, 33 H3K4me3). Model performance was high, with average area under the curves (AUCs) of 0.82–0.95 across assays (**Extended Data Fig. 8b**). Robustness was confirmed using OCR datasets from dairy cattle^27^ with a median AUC of 0.95.

We then scored 28 million SNPs derived from 3,530 whole genome sequences^55^ across diverse cattle breeds. We observed that deltaSVM scores were positively correlated with effect sizes of AS SNPs across epigenomic assays (**Fig. 5a, Supplementary Figure 28**), validating their use as proxies for regulatory impact of allelic changes. Following a strategy used in humans^18^, we then defined a “regulatory magnitude” for each SNP as its maximum deltaSVM score across all contexts. This score was weakly but significantly correlated with both context specificity (i.e., the number of contexts in which the SNP overlapped with regulatory peaks) and genomic proximity to TSS (i.e., the distance of a SNP to the nearest gene TSS) (**Fig. 5b-c**), as well as with conservation metrics (phyloP and GERP), but not with the FAETH score^56^. We further categorized SNPs by regulatory context overlap: 1) Null: no overlap; 2) Specific: overlapping peaks in a single context; 3) Multiple: overlapping peaks in more than one but fewer than 80% of contexts; and 4) Shared: overlapping peaks in more than 80% of contexts. Stratification of SNPs by regulatory overlap revealed that tissue-shared variants were enriched near TSS and associated with PLS, whereas tissue-specific variants overlapped primarily with ELS (**Fig. 5d-e**). Moreover, both eQTL and trait-associated QTLs were significantly enriched for variants with regulatory magnitude scores (**Supplementary Figure 28c**).

**Figure 5.**
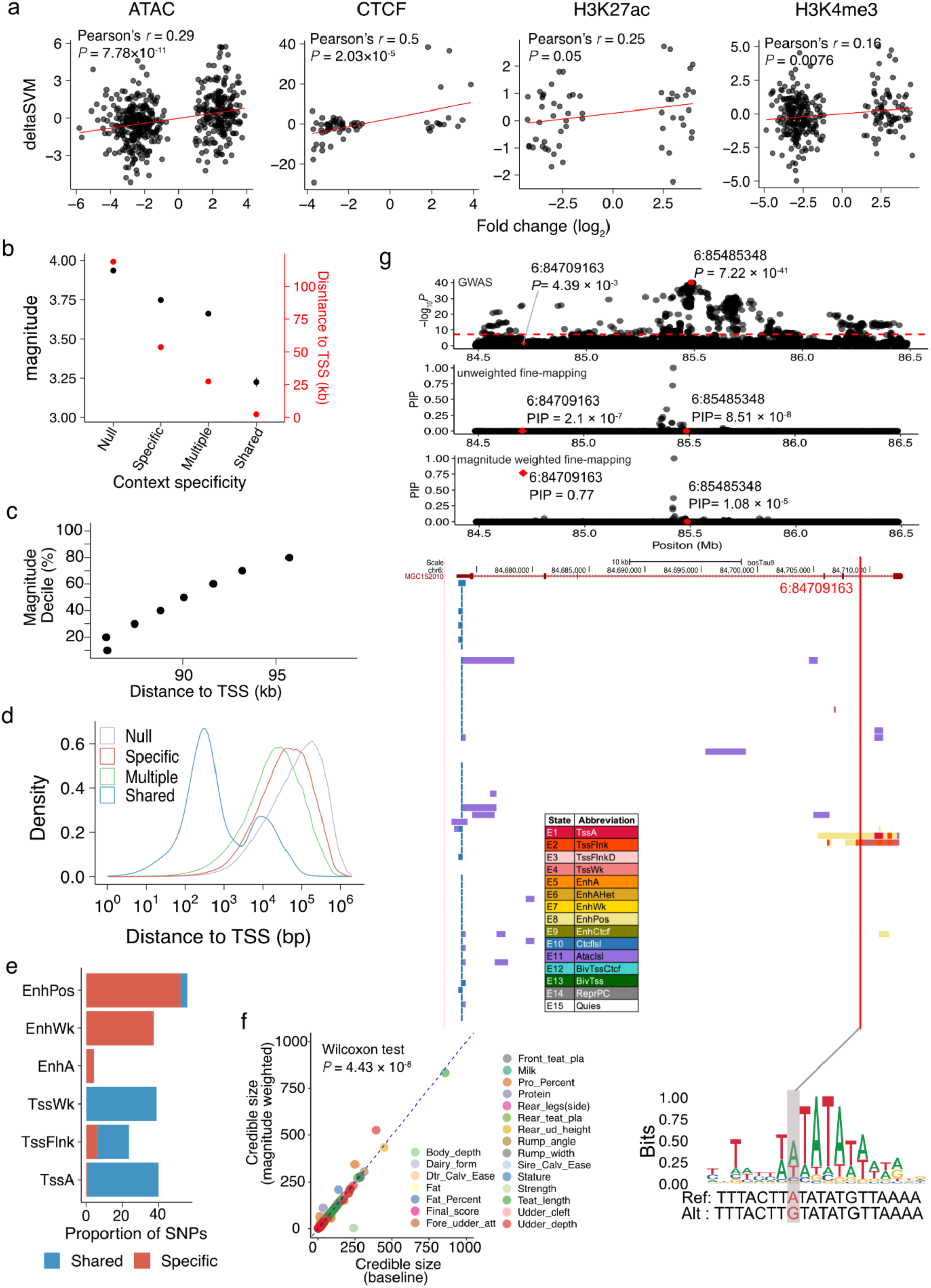
Predicting regulatory effects of sequence variants. (a) Correlation between deltaSVM scores (y-axis) and allelic fold change (log₂-scaled, x-axis) across four assays: ATAC-seq, CTCF, H3K27ac, and H3K4me3 (left to right, respectively). (b) Relationship between regulatory magnitude (left y-axis: defined as the maximum deltaSVM score across all assay-tissue contexts) and distance to the nearest transcription start site (TSS; right y-axis) as a function of context specificity (i.e., number of assay–tissue contexts affected). (c) Mean distance to TSS for genetic variants stratified by deciles of regulatory magnitude. (d) Density plot showing the distribution of variant distances to TSS categorized by context specificity, including: “null” (impacting 0 assay-tissue contexts), “specific” (impacting only 1 assay-tissue context), “multiple” (impacting > 1 but <80% of assay-tissue contexts); and “shared” (impacting >80% of assay-tissue contexts). (e) Proportion of SNPs residing within enhancer-like chromatin states (EnhPos, EnhWk, EnhA) and promoter-like chromatin states (TssWk, TssFlnk, TssA), stratified by “Specific” and “Shared” variant categories. (f) Regulatory magnitude weighted fine mapping produces smaller median credible set sizes across 22 complex traits (*P*=4.43 × 10^-8^, Wilcoxon test). (g) Fine-mapping a cattle GWAS locus for protein percentage in milk using regulatory magnitude as prior weights in SuSiE-adj^97,104^. The panels from top to down are: GWAS significance plot (-log_10_*P*); posterior inclusion probability (PIP) for fine-mapped variants without prior weights; posterior inclusion probability (PIP) for fine-mapped variants with regulatory magnitude prior weights; chromatin state annotations across 65 bovine epigenomes. The vertical red line highlights the fine-mapped variant overlapping both chromatin states and transcription factor binding sites (TFBS).

To evaluate their utility for fine mapping, we incorporated regulatory magnitude–based priors into Bayesian analyses of 22 complex traits (**Table S17**). This approach significantly reduced credible set sizes compared to unweighted analyses (**Fig. 5f**). For protein percentage, for example, regulatory weighting identified an additional putative causal SNP (6:84709163) (**Fig. 5g**). This variant is located in the intronic region of *MGC152010*, which is highly expression in liver (**Supplementary Figure 28d**, **Fig. 5g, Table S17**), overlaps a liver-specific enhancer, and is predicted to disrupt high-mobility group (HMG) factor binding (**Fig. 5g**). These results demonstrate that incorporating predicted regulatory effects into GWAS fine-mapping enhances causal-variant resolution, providing a framework for functional prioritization in livestock.

### Comparative analysis of bovine and human epigenomes

To explore the evolutionary conservation of *cis*-regulatory elements (CREs), we compared chromatin states between cattle and human across 22 matched tissues. For humans, we analyzed 1,089 Roadmap Epigenomics datasets^17^ spanning six regulatory marks (OCR, H3K27ac, H3K27me3, H3K4me1, H3K4me3, and CTCF). Human chromatin states were re-annotated using our pipeline to ensure methodological consistency, producing comparable emission profiles and state numbers in both species (**Fig. 6a**). We classified human CREs into four conservation categories: (1) sfCREs: conserved in both sequence and function; (2) sdCREs: conserved in sequence but divergent in function; (3) soCREs: conserved in sequence only; and (4) ssCREs: human-specific elements (**Fig. 6a**). Approximately 30% of human CREs were sdCREs, indicating dynamic functional divergence despite sequence conservation. PLS were more often functionally conserved (sfCREs) than ELS (**Fig. 6b**). Focusing on EnhA, sequence-conserved types (sfCREs, sdCREs, and soCREs) exhibited significantly higher phyloP scores than ssCREs (*q*-value < 10^-300^), with sfCREs being the most evolutionarily constrained (**Fig. 6c**). Notably, ssCREs were more proximal to TSS, whereas sequence-conserved elements, particularly sfCREs, were enriched at distal regions (**Fig. 6d**).

**Figure 6.**
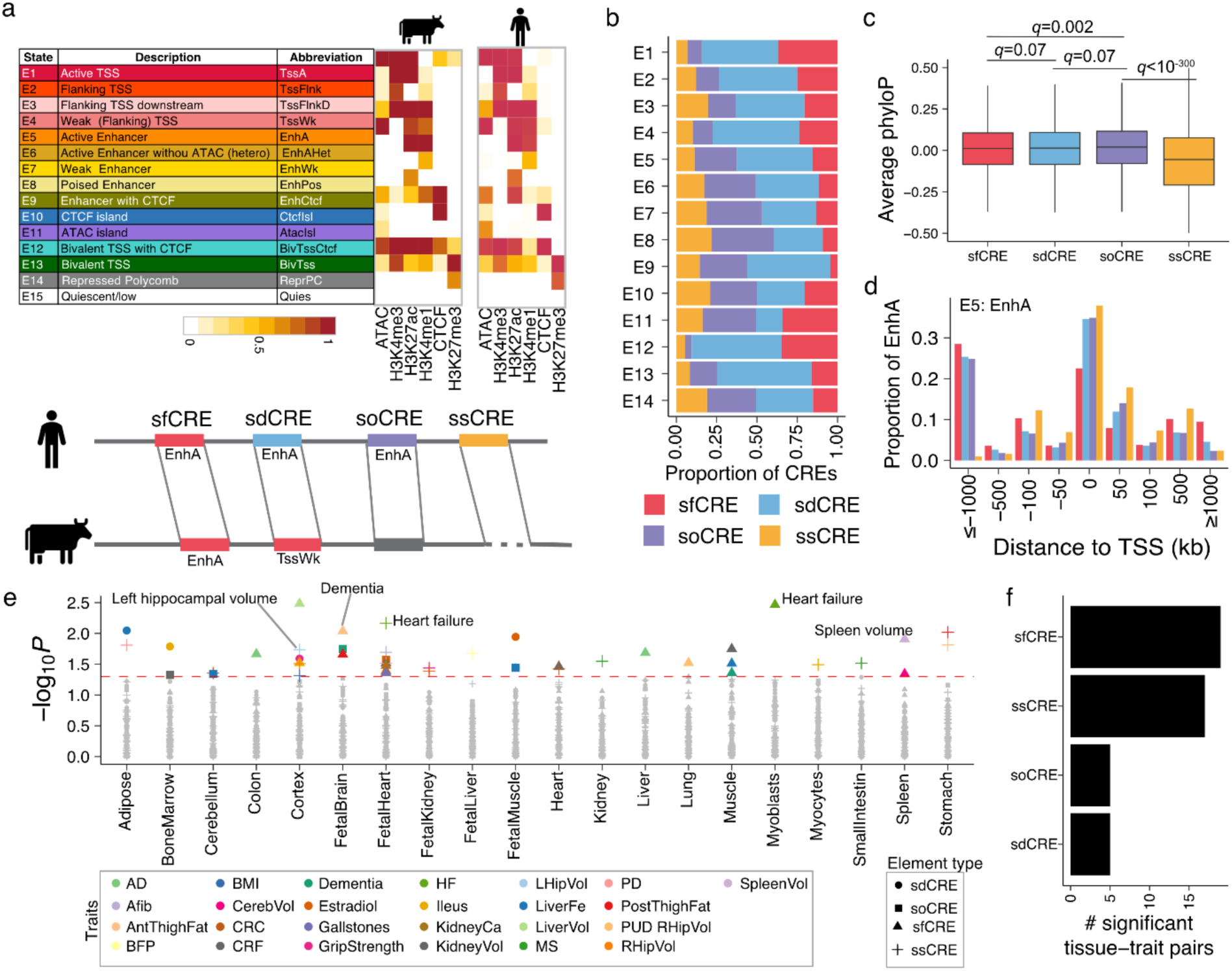
Cross-species comparison and conservation of regulatory elements. (a) Chromatin state annotations comprising 15 states were independently predicted in cattle and human. The top panel summarizes chromatin state labels, functional descriptions, abbreviations, and corresponding emission probabilities for both species. The bottom panel categorizes cis-regulatory elements (CREs) based on conservation: 1) sfCREs, sequence- and functionally conserved elements; 2) sdCREs, sequence conserved but functionally divergent; 3) soCREs, sequence-only conserved; and 4) ssCREs, human-specific elements. (b) Proportions of regulatory elements stratified by conservation category. (c) phyloP conservation scores for strong enhancers (EnhA), grouped by conservation category. Significance was assessed using a two-sided Student’s t-test with FDR adjustment. phyloP scores were obtained from Cactus 241-way mammalian alignments provided by the Zoonomia Project^105^. (d) Genomic distance between strong enhancers (EnhA) and the nearest transcription start site (TSS), stratified by conservation category. Color legend is same as panel (b). (e) Tissue-specific heritability enrichment of EnhA elements stratified by conservation category, represented as −log_10_*P* values. (f) Number of significantly enriched tissue– trait pairs (x-axis) contributed by EnhA elements from each conservation category (y-axis). Trait name abbreviations are provided at **Table S18.**

To assess trait relevance, we performed heritability partitioning using Linkage Disequilibrium Score Regression (LDSC)^57^ across 25 human GWAS traits (**Table S18**). The analysis identified 46 significant tissue-trait associations (FDR < 0.05), including enrichment of heart failure heritability in myoblasts and fetal heart (**Fig. 6e**). Remarkably, sfCREs and ssCREs contributed disproportionately to these enrichments, accounting for 41.3% and 37.0% of all significant tissue-trait pairs, respectively (**Fig. 6f**). These findings highlight that both evolutionarily conserved and lineage-specific enhancers play prominent roles in shaping the genetic architecture of human complex traits.

## Discussion

Here we present a comprehensive bovine multi-omic epigenomic atlas (1,138 datasets) that annotates approximately 45% of the genome as putative regulatory elements across tissues, developmental stages, and primary cell types. Beyond scale, the resource integrates chromatin states, accessibility, methylation, 3D contacts and gene expression to contextualize complex traits and extend prior coverage by ∼20%^11^. A key insight is strong context-specificity of enhancer activity and motif usage. The 10-state/15-state chromatin models reveal broad enhancer coverage with strong tissue specificity (notably in muscle, with fewer unique EnhA elements in digestive tissues), motif enrichments align with lineage programs (e.g., MYOG/MYF5/MYOD in muscle; NEUROD1/NEUROG2 in CNS), and thousands of tissue-specific enhancers cluster near canonical regulators (e.g., MYF5/MYF6). ABC links support a distributed, many-to-many enhancer–gene architecture (median ∼7 enhancers per gene and ∼7 genes per enhancer; mean distance ∼168 kb).

Leveraging this atlas, we applied S2F modeling that adds a mechanistic layer. ChromBPNet generalized well across 55 contexts and identified 301 motifs (with frequent KLF15-like motifs^47^), while deltaSVM-derived “regulatory magnitude” scores correlated with allele-specific effects and prioritized variants, improving the resolution of GWAS fine-mapping (credible-set reductions across 22 traits; illustrative new candidate at 6:84,709,163 near *MGC152010*). Integrating >91k allele-specific events with CattleQTLdb, identifies 439 putative functional variants that show assay- and tissue-matched effects (e.g., PKLR in jejunum; RTN3 in muscle), exemplifying a practical path from regulatory signal to candidate causal variants and tissues of action. Cross-species analyses indicate conserved promoters and lineage-specific enhancers both contribute to heritability, underscoring shared and divergent regulatory logic across mammals.

Despite these advances, several limitations remain. First, bulk tissue datasets limit resolution, masking cell-type–specific activity. Single-cell epigenomic and transcriptomic profiling is needed to capture cellular heterogeneity and dynamic transitions^58^. Second, the small cohorts limit detection of population-level regulatory variation. Population-scale efforts will enable molecular QTL discovery, gene regulatory network inference, and enhance precise genotype–phenotype links^10,19,21,50,59,60^. Third, greater sequencing depth and data quality will improve sensitivity for detecting weak or transient regulatory elements, and their annotation and enable assessment of complex variants^61–63^, such as structural variants^64,65^, which are currently underexplored in functional contexts. Finally, S2F models^66^ should be extended to multimodal, single-cell contexts, and paired with functional assays (e.g. MPRA^67^ and STARR-seq^68^) to establish causality.

In summary, by integrating chromatin state maps, enhancer–gene connectivity, sequence-inferred syntax, and complex trait integration, this atlas provides a practical foundation to interpret non-coding variation in cattle, accelerate hypothesis-driven validation in target tissues and traits, and accelerate precision breeding and comparative regulatory genomics.

## Acknowledgments

We thank all of the people that contributed to the breeding and collection of tissues used in this study. We also thank all the researchers who have contributed to the publicly available data used in this research. Stephanie D. McKay from University of Missouri for helping with the acquisition of the NIFA funding. Portions of this research were conducted with the advanced computing resources provided by Texas A&M High Performance Research Computing.

## Funding

This study was supported by Agriculture and Food Research Initiative Competitive Grant no. 2018-67015-27500 (H.Z., P.J.R. etc.) and sample collection was supported by no. 2015-67015-22940 (H.Z. and P.R.) and 2012-67015-23673 (MR) from the USDA National Institute of Food and Agriculture and OECD Co-operative Research Programme, 2012 Fellowship, TAD/PROG JA00074636 (MR). J.B. received scholarships financed by Foundation to Support the Research of São Paulo State (FAPESP) (Grants Numbers: 2020/01369-1 and 2022/12631-4). H.Z. acknowledges funding from the USDA National Institute of Food and Agriculture, Multistate Research Project NRSP8 and NC1170 and the California Agricultural Experimental Station. C.A.G. acknowledges support from Texas A&M AgriLife Research. B.M.M, acknowledges the support from the University of Idaho, the AVFS Meat Center, and the NRSP8 multistate Hatch Funding IDA 01566.

## Author contribution statement

H.Z., L.F. and D.G. conceived the manuscript. B.M.M., G.K.M., and K.D. collected the fetal tissues for all the molecular assays and K.D. performed the fetal ChIP assay. M.R. collected mammary gland tissues. D.G. conducted raw data preprocessing, peak calling, epigenome imputation, and chromatin state annotations. Z.J., W.L., G.R., and S.C. generated the RNA-seq data for all adult and fetal tissues. J.B. and D.G. analyzed RNA-seq and linked enhancer to target genes, J.B., D.G., C.A.G, G.W. and Y.X. processed WGBS data, D.G. and W.Y. annotated super-enhancers. W.Y., J.B., D.G., B.P., and C.P.W. performed allele-specific analysis, C.P.W., D.G. analyzed chromatin accessibility data, D.G., J.W. and J.J. conducted complex trait integration analysis, S.C., X.X., Y.Wang, Y. Z. and D.G. led new data generation and quality control for adult ChIP-seq data, H.L., H.Z. and L.F. provided eQTL resources used in this study, Y.U., V.M.R, and J.D. provide GWAS summary statistics from beef cattle, L.M., C.M., J.W. and J.J. provided GWAS summary statistics from dairy cattle. G.W. and Y.X. analyzed ATAC-Seq. B.M.M., K.D., P.L. and H.J. generated ChIP-seq data and primary cells. G.R., C.A.G., B.M.M, T.P.L.S., H.J., Z.J., W.L., J.R., J.F.M., J.D., P.J.R., J.E.K., M.R., H.Z. contributed to funding acquisition. D.G. prepared the initial manuscript draft with inputs from L.F. and H.Z. D.G, H.Z. and L.F. revised and finalized the manuscript. All authors read, edited and approved the final manuscript.

## Competing interests

Y.T.U., V.M.P.R. and N.D. are employed by STgenetics. P.J.R. acquired the funding for this research while affiliated to the University of California, Davis, and is currently employed by STgenetics. S.C. is from Zoetis Inc.

## Data and materials availability

We have deposited newly generated RNA-Seq, ChIP-Seq, ATAC-seq WGBS in FAANG portal (https://data.faang.org/), NCBI SRA database (https://www.ncbi.nlm.nih.gov/sra/), and European Nucleotide Archive. Accession IDs will be available upon the acceptance of the manuscript. Accession IDs for public data includes: PRJEB41939, PRJNA672996, PRJNA531208, E-MTAB-11825 and E-MTAB-11826. Metadata details are available in Supplementary Tables S1-S4. All processed data are available at Zenodo database (https://zenodo.org/10.5281/zenodo.12216791, available upon the acceptance). Cattle traits QTLs retrieved from CattleQTLdb (https://www.animalgenome.org/cgi-bin/QTLdb/BT/index), and eQTL data retrieved from https://cattlecell.kiz.ac.cn. Other data sets used in this study, published previously, are described in the Methods. All the computational scripts and codes are available at the GitHub website, including https://github.com/guandailu/BovineFAANG, https://github.com/guandailu/ChIPQC, and https://github.com/guandailu/SnakeHiC.

## Online Methods

### Sample collection

Samples used in this study were collected by the Functional Annotation of Animal Genomes (FAANG) community. The sampling procedure for adult tissues has been previously described in detail by Tixier-Boichard et al^69^. Briefly, tissue samples for beef cattle were collected from two male and two female L1 line Hereford cattle within 1–2 hours post-euthanasia, flash-frozen in liquid nitrogen, and stored at −80°C until further processing. For fetal tissues, four pregnant L1 Hereford were euthanized and the fetuses were collected at 78 days of gestation under the University of Idaho Institutional Animal Care and Use No. 2017-67. Fetal tissues were collected, flash frozen and stored at −80°C. Mamary gland tissues were collected by biopsy, using the Farr method^70^, at 4 time points mid-pregnant at day 100 of pregnancy, late pregnant ∼2 weeks pre-calving, early lactation ∼2 weeks post-calving, and adult state ∼36 month of age, tissue was flash frozen, see Beiki et al.^15^ for details. All animal experiments were conducted in accordance with Protocol for Animal Care and Use No. 18464, approved by the Institutional Animal Care and Use Committee at the University of California, Davis.

### Library preparation and sequencing

For RNA-seq experiments, frozen tissue samples were pulverized in liquid nitrogen using a mortar and pestle. Total RNA was then extracted with TRIzol reagent (Thermo Fisher Scientific, #15596026) and treated with DNase I (Thermo Fisher Scientific, #EN0521). RNA integrity was assessed using an Agilent Bioanalyzer (Agilent Technologies, Santa Clara, CA, USA), and samples with RNA integrity number (RIN) > 8 were used for library preparation, followed by sequencing on an Illumina Hiseq 4000 platform (Illumina, San Diego, CA, USA). For ChIP-seq (targeting H3K4me3, H3K4me1, H3K27ac, H3K27me3, and CTCF) experiments, samples were processed as previously described^71–73^. ATAC-seq libraries were prepared as described in refs^15,74,75^. Both ATAC-seq and ChIP-seq libraries were sequenced on an Illumina HiSeq 4000 platform using 50-bp paired-end and single-end reads, respectively. For whole-genome bisulfite sequencing (WGBS), genomic DNA of tissue samples was extracted, fragmented, and sequencing libraries were prepared using methylation sequencing adaptors. Libraries were then treated with sodium bisulfite, which converted unmethylated cytosines to uracils while preserving methylated cytosines. The bisulfite-converted libraries were then PCR-amplified and subjected to Illumina paired-end 150 bp sequencing to assess genome-wide DNA methylation at single-base resolution.

### Raw data processing and quality control

ChIP-seq sequencing data was processed using a uniform pipeline available at https://github.com/guandailu/ChIPQC, after gathering newly generated sequencing data in this study and previously published datasets (details provided in **Table S1**). Briefly, sequencing adapters were trimmed using Trim Galore (v0.6.7) (https://github.com/FelixKrueger/TrimGalore). Cleaned reads were aligned to the ARS_UCD1.2 reference genome (Ensembl release v105) using BWA-MEM^76^. For simplicity, we only considered the autosomal genome throughout this manuscript. PCR duplicated reads were removed prior to peak calling. Deduplicated BAM files were used to identify peaks with MACS2^77^ with the options of “--q 0.01” for narrow marks (i.e. H3K27ac, H3K4me3, CTCF), and of “--q 0.05 --broad” for broad marks (i.e. H3K4me1, H3K27me3, H3K36me3, and H3K9me3). Then, we filtered out samples with clean reads < 10 million, number of peaks < 10,000 and Jensen-Shannon Divergence (JSD) between two replicates < 0.05 for broad marks and < 0.1 for narrow marks. This filtering step finally left 607 datasets that remained for subsequent analyses. For ATAC-seq, we applied the same processing steps, including adaptor trimming, read alignments, the removal of PCR duplicated reads, and peak calling, while the peak calling was applied with the specific options of “-f BEDPE --nomodel”. The filtering criteria for ATAC-seq included a FRiP score >0.3 and checked TSS enrichment manually. For RNA-seq, a similar pipeline was applied while the read aligner STAR (v2.7.0)^78^ was used. Gene expression levels were quantified as Transcripts Per Million (TPM) using StringTie2^79^, and raw read counts were obtained using featureCounts (release 2.0.6)^74^ based on the ARS_UCD1.2 gene annotation (Ensembl release v105). 12 public Hi-C datasets were uniformly processed using the pipeline SnakeHiC (https://github.com/guandailu/SnakeHiC). For DNA methylation data generated by WGBS, adaptors were trimmed using Trim Galore (v0.6.6) and reads were preprocessed using the Bismark pipeline (Version 0.24.2). After further removing PCR duplicated reads, the alignment files in BAM format were subsequently converted into compressed formats, including “beta” and “pat”, using wgbs_tools (version 0.2.2)^80,81^. Homogeneously methylated blocks (defined as contiguous CpG sites with similar methylation levels within 5kb) were identified using the “*segment*” command, and blocks with fewer than 4 CpG sites were excluded from downstream analyses. For each segmented block, we computed the average methylation level across CpG sites based on the corresponding *beta* formatted files.

### Imputation of missing and low-quality histone and CTCF signals

To obtain all the core histones marks (i.e., H3K4me1, H3K4me3, H3K27ac, and H3K27me3) and CTCF for chromatin state annotations, we used the ChromImpute pipeline (v1.0.5) developed by the Roadmap Epigenomics Consortium^17,29,30^. Briefly, we generated signal tracks for experimental samples using the *bdgcmp* function of the MACS2 tool^77^, computing −log_10_*P* as recommended by Kundaje et al.^29^ and Ernst et al.^30^. For brevity, we only considered signals for autosomes 1-29. ChIP-seq signal tracks were converted to 25-bp resolution by averaging signal intensity across each 25-bp bin. Imputation of missing or low-quality signals followed four main steps: 1) computing global distance between datasets with the *ComputeGlobalDist* function; 2) generating training features with the *GenerateTrainData* command; 3) training mark-specific and sample-specific predictors with *Train* command; and 4) applying these predictors to generate imputed signal tracks using *Apply*. To assess imputation accuracy, we implemented a cross-validation-like procedure. Two experimental samples were randomly masked, and imputation was performed on these held-out samples. The correlation between imputed and observed signals was then assessed using the *Eval* function. This process was repeated 15 times to ensure that each epigenetic mark was tested at least twice. In addition, imputed signals were manually inspected to ensure quality. Additionally, we validated the robustness of the pipeline using ChIP-seq datasets published by Prowse-Wilkins et al.^28^, masking 15 samples using the same procedure. Comparable high imputation accuracy was observed across these held-out datasets.

### Predicting chromatin states and super-enhancers

We used ChromHMM (v1.26)^31,32^ to predict chromatin states for 72 bovine epigenomes, each of which included the core set of four histone modifications: H3K4me1, H3K4me3, H3K27ac, and H3K27me3. Briefly, we used the *LearnModel* function in ChromHMM to train models with state numbers ranging from 2 to 15—the maximum number supported by the software. To identify the optimal model, we compared emission probability across models following the approach described by Gorkin et al.^82^ The optimal number of states was determined as the point at which the median correlation of emission probability across two or more replicates plateaued. For the 72 bovine epigenomes, a 10-state model was ultimately selected, and state labels were manually assigned based on an extensive literature review, particularly as described in refs^31–33^. For 65 bovine tissues with core epigenome histones with two additional marks (OCRs and CTCF), we ran a similar pipeline and learned 15 chromatin states. In order to compare cattle data with that of humans, we also conducted the chromatin state annotation for 134 human epigenomes with the same pipeline.

We also annotated super-enhancers, a class of regulatory elements that differ from typical enhancers in terms of their size, transcription factor density and content, transcriptional activation potential, and sensitivity to perturbation^35,83^. Super-enhancer identification was performed using the ROSE algorithm (version 0.1)^35^ across 33 tissue contexts. For each tissue, H3K27ac ChIP-seq profiles from two biological replicates were merged using SAMtools (v1.19.2)^84^. As input, we used the genomic regions annotated as typical enhancers (i.e. chromatin states E4, E5, and E6 from 10-state model) to define candidate super-enhancer regions. These were then ranked based on H3K27ac signal intensity to identify super-enhancers following the standard ROSE pipeline^35^.

### Linking enhancers to genes

To predict target genes of enhancer elements, we employed the Activity-by-Contact (ABC) model^34^. We applied the ABC pipeline across 62 epigenomes from 37 tissues that had matched ATAC-seq and H3K27ac ChIP-seq data. The ABC score for a given enhancer-gene pair was calculated as:

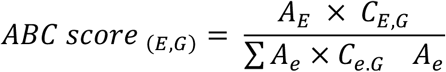

where E and G represent the enhancer element and the gene, respectively. A(E) represents the activity of element E, and C(E, G) denotes the Hi-C contact frequency between element E and the promoter of gene G. The denominator sums this product over all candidate elements within ±5 Mb of the gene. Activity was quantified using read counts from ATAC-seq and H3K27ac ChIP-seq data. Candidate enhancer elements were defined as ∼500 bp regions centered on ATAC-seq peaks.

### Context specificity analyses

#### For gene expression

We employed the Tspex^85^ as described previously^59^. For a given tissue, we computed the average gene expression across biological replicates. Tissue-specific expression was then defined using a Z-score approach across all tissues, with genes considered tissue-specific if Z-score >= 0.8. Adult and fetal tissues were separate in this analysis. Subsequently, we performed enrichment analysis on the identified tissue-specific gene sets using the Enrichr package (v3.2)^86^ and the GO_Biological_Process_2023 database. The significant threshold was set as adjusted *P*-value threshold of <= 0.05 with the False Discovery Rate (FDR) approach^48^.

#### For chromatin states

We applied an intuitive comparison between the target tissue and the test to identify tissue-specific chromatin states with BEDtools (version 2.31.0)^87^. That is, a state observed in one tissue but not in others. Motif enrichment of tissue-specific chromatin states was conducted using the Hypergeometric Optimization of Motif EnRichment (HOMER) tool (v5.1, 7-16-2024)^36^. *For open chromatin regions*: We identified differentially accessible peaks by gathering all OCR peaks from valid samples to create the consensus set of peaks. Then we quantified raw reads counts of consensus peaks across samples with featureCounts (v1.6.2)^88^. Through comparing a given tissue with the rest using edgeR tool (version 4.4.0)^89^, differentially accessible peaks were identified, which were deemed as tissue-specific OCRs. The significant threshold was set as FDR-adjusted *P* < 0.05 and fold change at log_2_ scale > 0.

#### For DNA methylation

Tissue-specific differentially methylated regions (DMRs) were identified by comparing each focal tissue against all others using the *find_markers* function in the wgbs_tools^80^ package (v0.2.2). Statistical significance was assessed via two-sided t-tests, and DMRs with *P* > 0.05 were excluded using the --pval 0.05 option.

#### Comparing adult and fetal samples

We identified differential chromatin states between adult and fetal stages using an BEDtools (version 2.31.0)^87^. That is, a state observed in one stage but not in the counterpart.

### Allele-specific identification

We identified genome-wide single nucleotide polymorphisms (SNPs) from two biological replicates (M08 and M22, two males) using the Genome Analysis Toolkit (GATK, v4.5.0.0) following the “best practices” pipeline^90^. To minimize reference mapping bias and enable accurate detection of allele-specific (AS) events, we applied the WASP (v0.16.3) pipeline^91^ to all epigenomic datasets. Read counts for each SNP were quantified using the *ASEReadCounter* module in GATK. SNPs were retained for AS analysis if they (i) were heterozygous, (ii) overlapped epigenetic peaks previously identified by MACS2^77^, (iii) were located on autosomes, and (iv) had >6 total reads, with each allele supported by at least 3 reads. For each retained SNP within a sample, a two-sided binomial test was performed to assess allelic imbalance. *P-*values were then adjusted for multiple testing using the Benjamini-Hochberg (BH) procedure^48^, and SNPs with a false discovery rate (FDR) < 5% were considered significant AS events.

### Training ChromBPNet to learn sequence motifs

We employed ChromBPNet (v1.0)^43^, a CNN-based deep learning framework, to learn sequence regulatory syntax of chromatin accessibility in the bovine genome. Briefly, for each of the 55 tissue contexts, we merged BAM files from biological replicates to generate input data. The analysis was restricted to ATAC-seq data, as ChromBPNet is specifically designed to model chromatin accessibility and also for simplicity. OCR peaks, previously called by MACS2^77^, were merged by tissue using the Irreproducible Discovery Rate (IDR) framework (version 2.0.3)^92^ and filtered to exclude regions overlapping with highly repetitive sequences. Genome mappability for the ARS-UCD1.2 reference genome was computed using GenMap (v1.3.0)^93^. ChromBPNet models were trained using a chromosome-based data split: training set (chr2, chr4-5, chr7, chr9-19, chr21-29), validation set (chr8, chr20), and testing set (chr1, chr3, chr6). As negative sets, we generated GC-matched background regions using the “prep nonpeaks” function of the ChromBPNet pipeline. Model training proceeded in two steps: (1) training a bias model to account for Tn5 insertion bias, and (2) training a bias-factorized ChromBPNet model. Chromatin accessibility in OCRs from the test set was then predicted using the *pred_bw* function, which outputs both read counts and signal shape of profiles. Prediction accuracy was evaluated using Pearson’s correlation, Spearman’s correlation, and JSD. In accordance with the original publication^43^, Pearson’s correlation between observed and predicted read counts demonstrated desirable evaluations and was therefore selected for subsequent analyses. To interpret model predictions and identify sequence motifs underlying OCRs, nucleotide-level contribution scores were computed and stored as BigWig files. These scores were used for *de novo* motif discovery with TF-MoDISCo (v1.0.0)^46^. To construct a non-redundant motif catalog across tissue contexts, we clustered predicted motif instances and merged clusters with pairwise similarity scores greater than 0.1, yielding a final set of 301 non-redundant sequence motifs.

### Training gkm-SVM classifier

To assess the regulatory impact of DNA sequence variants in the cattle genome, we adopted the deltaSVM approach^54^, which offers both high predictive accuracy and computational efficiency, making it suitable for genome-wide variant scoring across diverse epigenetic marks and tissue contexts. Specifically, we trained gapped k-mer Support Vector Machine (gkm-SVM) classifiers^94,95^ for each tissue-epigenetic mark combination. For each epigenetic mark, we first merged MACS2^77^ identified peaks from two biological replicates using the IDR framework (v2.0.3)^92^ and filtered out regions overlapping with highly repetitive sequences. The remaining peaks were then aggregated across all tissues to define a positive set. To generate the negative set, we sampled GC-matched genomic regions that did not overlap any peak regions. We used LS-GKM (v0.1.1)^96^, an optimized implementation of gkm-SVM with improved performance^95,96^, to train the classifiers. Model performance was evaluated using 10-fold cross-validation, with parallelization enabled via the “-i” option.

To compute deltaSVM scores, we first generated all non-redundant 11-mers using the *nrkmers.py* script. Then, for ∼28 million SNPs identified from 3,530 whole-genome sequences^55^, we synthesized 40-bp allelic oligonucleotides for both reference and alternative alleles (centered on the SNP with ±19 bp flanking sequence). Variant impact scores were computed using the deltasvm.pl script with the following command: perl deltasvm.pl chr${chr}.ref.fa chr${chr}.alt.fa weights.txt deltaSVM.tsv. For each SNP, we defined the maximum deltaSVM score across all tissue-mark combinatory contexts as its regulatory magnitude, representing the predicted change in regulatory activity between alleles. To assess whether CattleGTEx eQTLs^21,50^ and traits QTLs^5^ are enriched for high regulatory magnitude scores, we conducted an enrichment analysis by fitting a linear model. For eQTLs, we compared fine-mapped causal variants (posterior inclusion probability [PIP] > 0.8) to non-causal eQTL SNPs (PIP < 0.01). For trait QTLs, we contrasted them with randomly selected SNPs matched for minor allele frequency (MAF). In both analyses, we regressed out genomic confounders, including distance to the nearest TSS, context specificity, and GERP conservation scores, as these features were significantly correlated with regulatory magnitude.

### GWAS fine-mapping analyses

Genome-wide significant association peaks for 22 dairy bull traits were identified from GWAS summary statistics (Wang et al., in press) using a significance threshold of *P* < 5 × 10^-8^. For fine-mapping analysis, candidate regions were defined as 2-Mb windows centered on the lead SNP of each associated peak. Fine-mapping was performed using SuSiE-adj^95^, an adaptation of SuSiE^97^ that improves fine-mapping accuracy in samples with family relatedness by incorporating linear mixed model (LMM)-derived summary statistics. We applied SuSiE-adj both with and without scaled regulatory magnitude as prior weights across 22 traits. Conditional analysis was conducted by including the lead SNP as a covariate and performing association analysis using the SLEMM-GWA method^98^.

### Linkage disequilibrium score regression analysis

To investigate whether the conservation of regulatory elements defined in this study could inform the genetic architecture of human complex traits, we collected GWAS summary statistics for 25 traits (details in **Table S18**). We performed tissue-specific heritability enrichment analyses using the LDSC package (v1.0.0)^99,100^. GWAS summary statistics were standardized using the munge_sumstats.py script to remove SNPs with MAF ≤ 0.01, INFO score ≤ 0.9, genotype call rate ≤ 0.75, duplicate rsIDs, out-of-bounds *P*-values, extreme chi-squared statistics, strand ambiguity, or discordance with SNPs used in LD score calculations. For each category of regulatory elements, we identified tissue-specific segments by contrasting the target tissue with others using BEDtools (v2.31.0)^87^. Annotations were generated for LDSC analysis using the *make_annot.py* script, and stratified LD scores were then calculated within a 500 kb window using SNPs from the 1000 Genomes Project Phase 3. We used default SNP weights and reference panels obtained from https://zenodo.org/records/7768714. Tissue-specific heritability enrichment analyses were then conducted using the “--h2-cts” option in LDSC^98^.

## Notes

### Summary of Updates

This revision includes: 1) authorship; 2) results; 3) funding acknowledgment; 4) author contribution statement

